# Neurotransmitter signaling specifies sweat gland stem cell fate through SLN-mediated intracellular calcium regulation

**DOI:** 10.1101/2023.09.10.557066

**Authors:** Juliana Remark, Jie Tong, Meng Ju Lin, Axel Concepcion, Satvik Mareedu, Gopal J Babu, Stefan Feske, Catherine P. Lu

## Abstract

Sympathetic nerves co-develop with their target organs and release neurotransmitters to stimulate their functions after maturation. Here, we provide the molecular mechanism that during sweat gland morphogenesis, neurotransmitters released from sympathetic nerves act first to promote sweat duct elongation via norepinephrine and followed by acetylcholine to specify sweat gland stem cell fate, which matches the sequence of neurotransmitter switch. Without neuronal signals during development, the basal cells switch to exhibit suprabasal (luminal) cell features. Sarcolipin (SLN), a key regulator of sarcoendoplasmic reticulum (SR) Ca^2+^-ATPase (SERCA), expression is significantly down-regulated in the sweat gland myoepithelial cells upon denervation. Loss of SLN in sweat gland myoepithelial cells leads to decreased intracellular Ca^2+^ over time in response to ACh stimulation, as well as upregulation of luminal cell features. In cell culture experiments, we showed that contrary to the paradigm that elevation of Ca^2+^ promote epidermal differentiation, specification of the glandular myoepithelial (basal) cells requires high Ca^2+^ while lowering Ca^2+^ level promotes luminal (suprabasal) cell fate. Our work highlights that neuronal signals not only act transiently for mature sweat glands to function, but also exert long-term impact on glandular stem cell specification through regulating intracellular Ca^2+^ dynamics.

## Introduction

Autonomic nerves are essential for all vital physiologic functions of human body. The parasympathetic and sympathetic neurotransmitters (NT), acetylcholine and norepinephrine respectively, engage their corresponding receptors on target organs to regulate their functions, such as heartbeat, blood pressure, muscle movements, digestion, glandular secretion (1). It is known that NT signals from these autonomic nerves not only initiate rapid and transient responses in the mature target organs, but may also influence their morphogenesis and regeneration (2-8). In salivary glands, it was demonstrated that the progenitor cell population is maintained in an undifferentiated state upon the reception of acetylcholine from parasympathetic nerves and that restoration of parasympathetic function can also increase salivary gland regeneration after damage (4, 5). However, the molecular mechanisms underlying their long-term impact has not been elucidated.

Sweating, crucial for human thermoregulation and water balance, relies entirely on the neuronal stimuli. The sympathetic nerves innervating sweat glands were classically described as one of the unique cases that undergo a “neurotransmitter switch” from initially releasing sympathetic norepinephrine (NE) to parasympathetic acetylcholine (ACh) during development (9, 10); and further, this switch was shown to be target (sweat gland)-dependent: in tabby mice, which fail to develop sweat glands, sympathetic nerves could still grow into the region, but they fail to acquire cholinergic properties and later disappear (11). In addition, grafting developing sweat glands to another skin region can trigger the local sympathetic nerves switch to produce ACh, which otherwise only produce NE (12-13). Characterization of NT receptors in the sweat glands was done decades ago using various selective antagonists for receptors with poor resolution, which adds to the confusion and complexity of spatiotemporal NT reception by the developing sweat duct progenitors and their respective roles in sweat gland morphogenesis and functioning.

Sweat gland morphogenesis is a two-step process: the straight duct elongates first, then the cells at the tip of the duct differentiate into coiled glandular structures and grow until fully mature (14). Previous work had demonstrated the progenitor populations in the sweat ducts and glands exhibit distinct transcriptional profiles and have respective roles in injury repair and maintenance (15). Additionally, purified myoepithelial cells in the sweat glands were shown to have the ability to give rise to a sweat gland organoid upon engraftment, presenting a great potential for future *de novo* sweat gland regeneration (15). Nevertheless, to generate a fully functional sweat glands for future therapeutic purposes, it is critical to understand how and which neurotransmitter(s) from sympathetic nerves impact on sweat gland morphogenesis during the nerve-gland co-development.

In this work, we sought to understand the molecular mechanisms by which neural signals impact sweat gland development. Through denervation and knocking out specific NT receptor expression during sweat gland development, we demonstrated that the basal cells lost their cell fate identity by gaining the suprabasal (luminal) cell secretory machinery upon loss of neuronal signals. Further, we identified that sarcolipin (SLN), a key regulator of SERCA, is significantly down-regulated in sweat gland myoepithelial stem cells upon denervation, and that SLN knockout sweat glands exhibit defects in intracellular Ca^2+^ regulation and cell fate specification at both transcriptional and epigenetic level. Our work provides novel insights into the connections between neurotransmitters, intracellular calcium regulation, and cell fate specification critical for proper glandular development.

## Results

### Denervation causes defect in sweat gland stem cell maturation

To investigate the involvement of neuronal signals in sweat gland development, we ablated sympathetic nerves that innervate the sweat glands using 6-hydroxydopamine (6-OHDA) to treat early postnatal mice (P0-P5) when their sweat ducts start to develop (19). These mice were not able to sweat as adults (**Figure 1A**) and sympathetic innervation (tyrosine hydroxylase, TH+) did not appear around adult sweat glands (**Figure 1B**) indicating that the effects of early denervation last long-term (**Figure S1A**). We employed FACS to purify glandular basal (myoepithelial) cells (Itgα6^hi^ Sca1^neg^ Itgβ1^hi^) and ductal basal cells (Itgα6^hi^ Sca1^pos^ Itgβ1^lo^) from control and denervated adult foot skin and performed colony formation assay to examine their ability to form holoclones in culture. Both basal cell types from denervated mice exhibited a decreased efficiency in colony formation, suggesting that denervation during early morphogenesis causes long-lasting defects in proliferation (**Figure 1C-D**).

**Figure 1.**
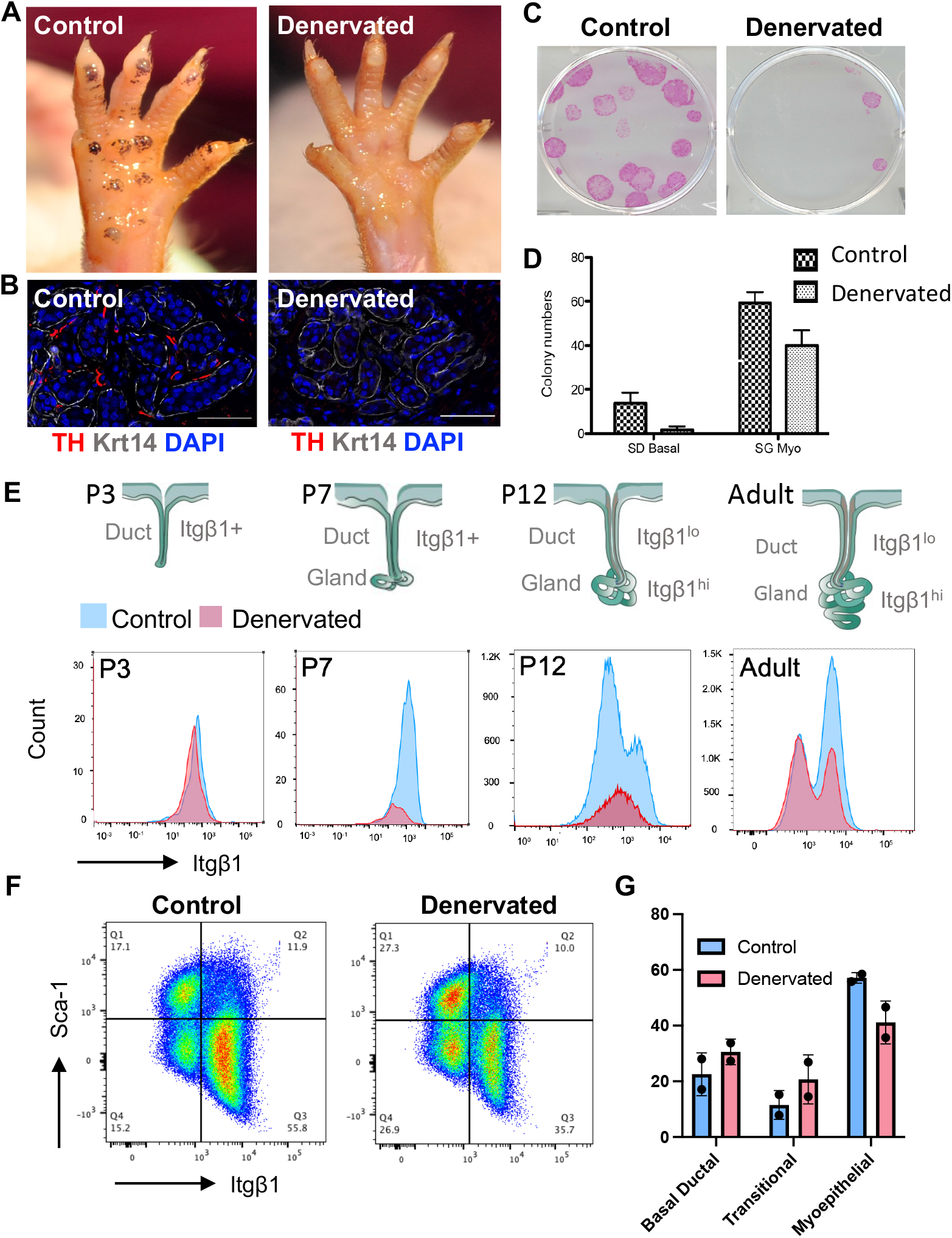
Denervation causes defects in sweat gland stem cell maturation. **(A)** Iodine-starch sweat test of natural sweating response after 3 min. Black dots indicate activate sweat pores; **(B)** Immunofluorescent microscopy with adrenergic nerve marker tyrosine hydroxylase (TH) in adult WT and denervated sweat glands marked by keratin 14 (Krt14); **(C)** Colony formation assay and **(D)** quantification; **(E)** Schematic showing sweat duct and gland development at various post-natal (P) stages (day 3, 7, 12), and adult. Integrin β1 (Itgβ1) expression, hi = high, lo= low, with representative FACS analysis histogram of Itgβ1 expression in basal cell keratinocyte populations during development below in control and denervated mice; **(F)** Representative FACS plots for basal cells from control and denervated mouse foot pads. **(G)** Quantification of FACS analysis of cell populations present n=2 with 3 mice used in each sort per condition. Basal ductal= Sca-1^hi^ Itgβ1^lo^, Transitional= Sca-1^lo^ Itgβ1^lo^, Myoepithelial= Sca-1^lo^ Itgβ1^hi^, ns between groups.

Further, we showed that as the duct invaginates into the dermis, the ductal progenitor cells down-regulate Sca1 and up-regulate Itgβ1 as they grow into the glandular region (**Figure S1B**). We used FACS analyses to monitor this glandular maturation process in control and denervated foot skin and found that although there is little difference in the expression of Itgβ1 between control and denervated samples at P3 as only the duct is formed at this point, defects in Itgβ1 upregulation start to show at P7 when glandular differentiation begins to take place. The segregation of ductal and glandular cells starts at P12 as shown by two distinct populations in the adult in control, but not in denervated foot skin (**Figure 1E)**. We noticed that during the maturation process, there is a population that contains Sca1^neg^ Itgβ1^lo^ cells, suggesting that they may be in transition from ductal cells to glandular cells. Importantly, we found that the denervated samples shows a higher percentage of ductal and transitional cells and a lower percentage of myoepithelial cells than the control samples (**Figure 1F-G**), together supporting the notion that the presence of nerves during development is critical for glandular maturation.

### Specific neuronal reception to NE vs ACh in sweat ducts and glands

It has long been established that the sympathetic nerves innervating sweat glands undergo a “neurotransmitter (NT) switch” from initially releasing the adrenergic NT norepinephrine (NE) to releasing the parasympathetic NT acetylcholine (ACh) later in development (10-11). However, it is unclear whether and how sweat ducts and sweat glands respond to both or either NTs and what their respective roles in glandular morphogenesis are. We employed single-cell RNA sequencing to acquire transcription profiles of wild-type adult and developmental (P0, P3, P6) mouse foot skin (**Figure S2A**). Using the signature genes that were previously identified from bulk RNAseq (15), we annotated the clusters representing different cell types in the sweat duct and gland (**Figure 2A-B, S2F**). Amongst all the NT receptors, we found that the Adrb2 is primarily expressed in the duct **(Figure 2C, E)** while the Chrm3 is primarily expressed in the gland (**Figure 2D, F**). Intriguingly, we found that the sweat gland myoepithelial cells have much higher expression of the Chrm3 receptor than the luminal cells **(Figure S2E)**, which are previously considered to be the main cell type responding to acetylcholine as they perform secretory functions (16).

**Figure 2.**
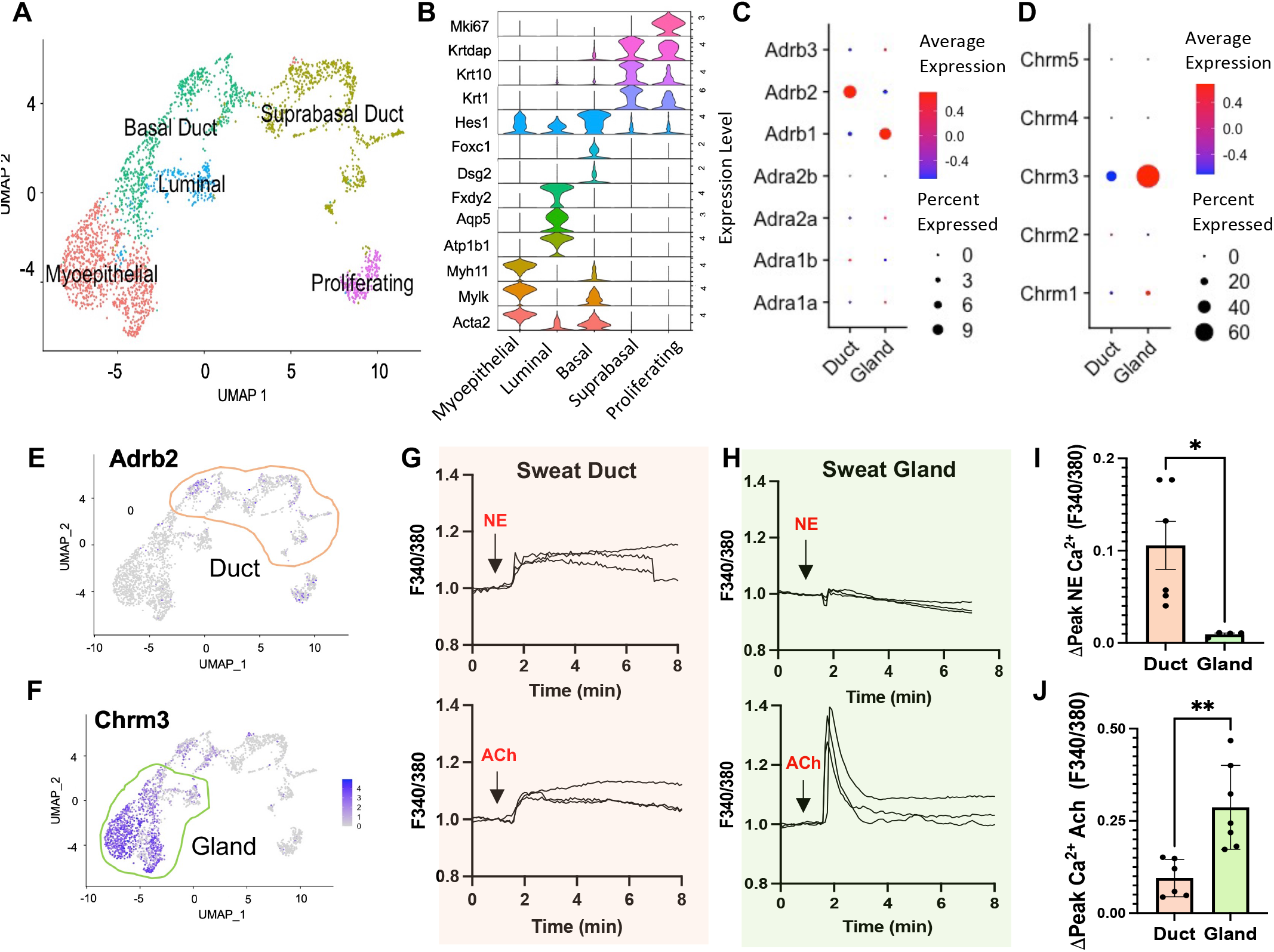
Specific neuronal reception to NE vs ACh in sweat ducts and glands. **(A)** UMAP projection of single-cell RNA-seq (scRNA-seq) data from combined sweat glands cells of P0, P3, P6, and adult mice **(B)** Violin plot showing the identity of sweat gland cell types was determined by established gene markers. **(C)** Dot plot of the expression of adrenergic NE receptors and **(D)** cholinergic ACh receptors in developing and adult sweat duct (suprabasal and basal duct populations) and gland (luminal and myoepithelial populations). Size of dot indicates percentage of expression within cells. Red indicates high expression while blue indicates low or no expression **(E)** Feature plot of Adrb2 expression and **(F)** Chrm3 expression. **(G)** Representative traces of the intracellular calcium response from isolated sweat ducts and **(H)** glands to NE (norepinephrine) and ACh (acetylcholine) loaded with the calcium indicator Fura-2. **(I)** Quantification of the peak response to NE (highest value after stimulus addition-baseline calcium before stimulus) in duct and gland, p=0.0183 **(J)** and peak calcium response to ACh in sweat glands and ducts, each dot represents an individual sweat gland or duct from 3 mice, unpaired t-test, p=0.0029

To examine the functionality of these receptors, we performed Ca^2+^ imaging using the calcium indicator Fura-2 for the isolated individual glands *ex vivo* and analyzed their responses to NE and ACh in the duct and gland. As shown in **Figure 2G-H**, the sweat ducts exhibit higher responsiveness to NE than ACh, while the glands exhibit responsiveness to ACh and not NE **(Figure 2I-J)**, validating the expression of functional receptors for NE in the sweat duct and for ACh in the sweat gland. Collectively, these results show that the sweat duct and gland have distinct responses to the two autonomic neurotransmitters, NE and ACh, respectively.

### Adrenergic and cholinergic reception are important for ductal vs glandular fate

We sought to understand what role NE has in ductal developmental by generating mice with a skin-specific deletion of Adrb2 using K14Cre. With tissue whole mount staining and clearing, we acquired 3D-reconstructed images to capture the entire sweat apparatus in the mouse foot pads to precisely quantify the length of the sweat ducts and the size of the sweat glands. We found that the deletion of the Adrb2 receptor results in significantly shorter sweat ducts (**Figure 3A-B**). Further, by FACS analyses, we showed that there is a lower percentage of ductal cells compared to wild type control (**Figure 3C-D**), suggesting that NE reception is critical for ductal cell specification, and that ductal progenitor cells may proceed to develop into glandular formation in the absence of NE signaling.

**Figure 3.**
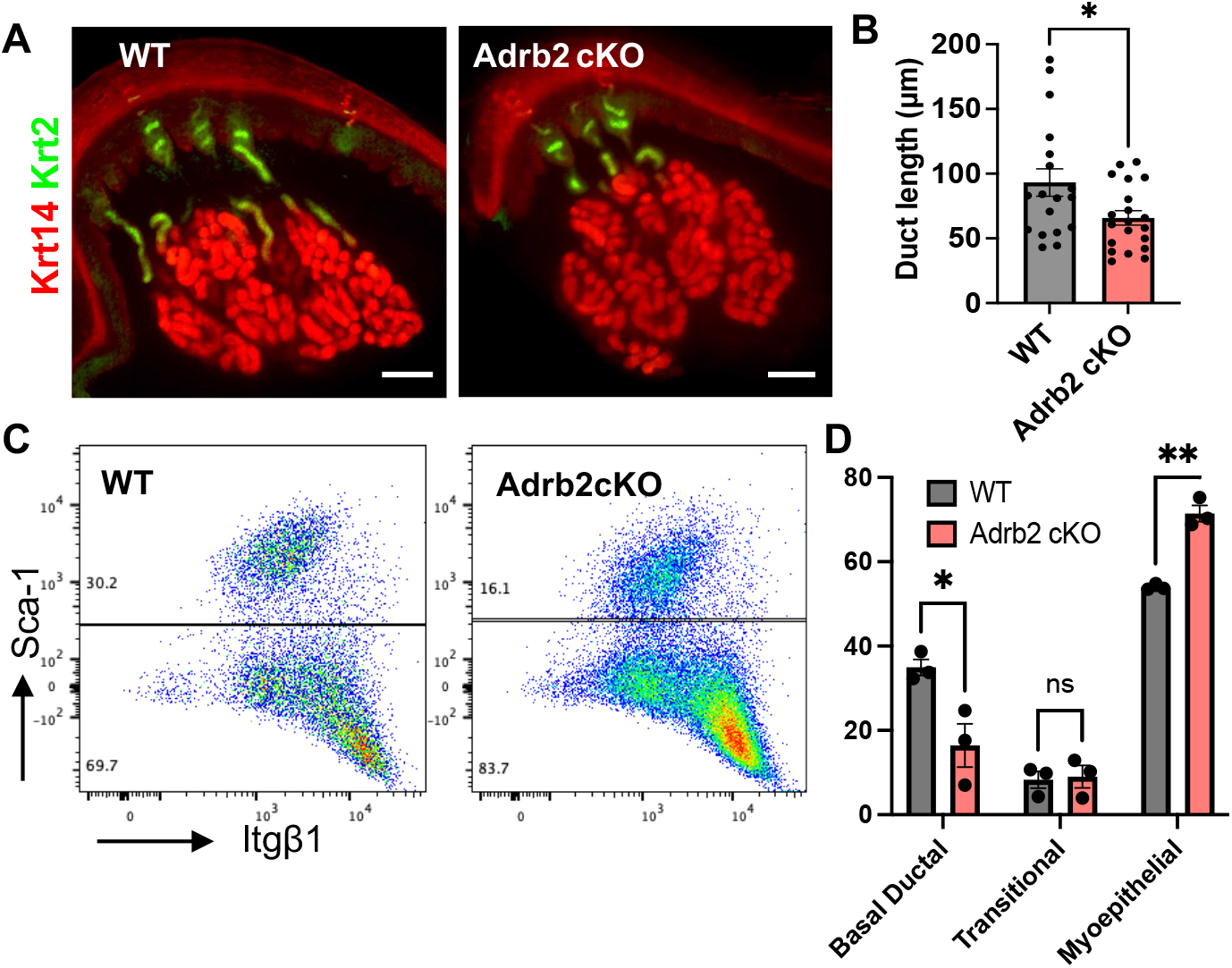
Adrenergic reception is important for ductal fate. **(A)** Representative slice from immunofluorescent image from whole-mount light-sheet imaging of WT and Adrb2 cKO sweat glands; keratin 14 (Krt14) keratinocyte marker, keratin 2 (Krt2) ductal marker **(B)** Quantification of duct length, unpaired t-test, p=0.0244, 6 mice used per condition, each dot represents a measured ductal length **(C)** Representative FACS plot of basal sweat gland populations in WT and Adrb2 cKO mice **(D)** Quantification of FACS analysis of cell populations present, n=3 with 3 mice used in each sort per condition, unpaired t-test, p= 0.029, p=0.028, Basal ductal= Sca-1^hi^ Itgβ1^lo^, Transitional= Sca-1^lo^ Itgβ1^lo^, Myoepithelial= Sca-1^lo^ Itgβ1^hi^

Next, using mice with a skin-specific deletion of the Chrm3 using K14Cre to examine the role of ACh signaling in glandular development, we found that Chrm3 cKOs grow significantly longer ducts **(Figure 4A-B)**. By FACS analyses, we showed that Chrm3 cKO exhibit a higher percentage of ductal and transitional cells, and a lower percentage of myoepithelial cells, indicating that ACh signaling through Chrm3 is important for glandular specification, and that without it, the progenitor cells fail to upregulate Itgβ1 and complete glandular maturation (**Figure 4C-D**). Note that this pattern of changes in cell type percentages is similar as shown in the denervated sweat glands (**Figure 1F-G**), indicating that neuronal signaling that impacts glandular development is through ACh acting on cholinergic receptors. Further, we analyzed the Ca^2+^ influx in response to ACh in wildtype control and Chrm3 cKO. We found that there is a lower elevation of intracellular Ca^2+^ level right after ACh stimulation, as well as overall lower Ca^2+^ level over time, albeit the Ca^2+^ response in not completely abolished in Chrm3 cKO sweat glands (**Figure 4E-G**).

**Figure 4.**
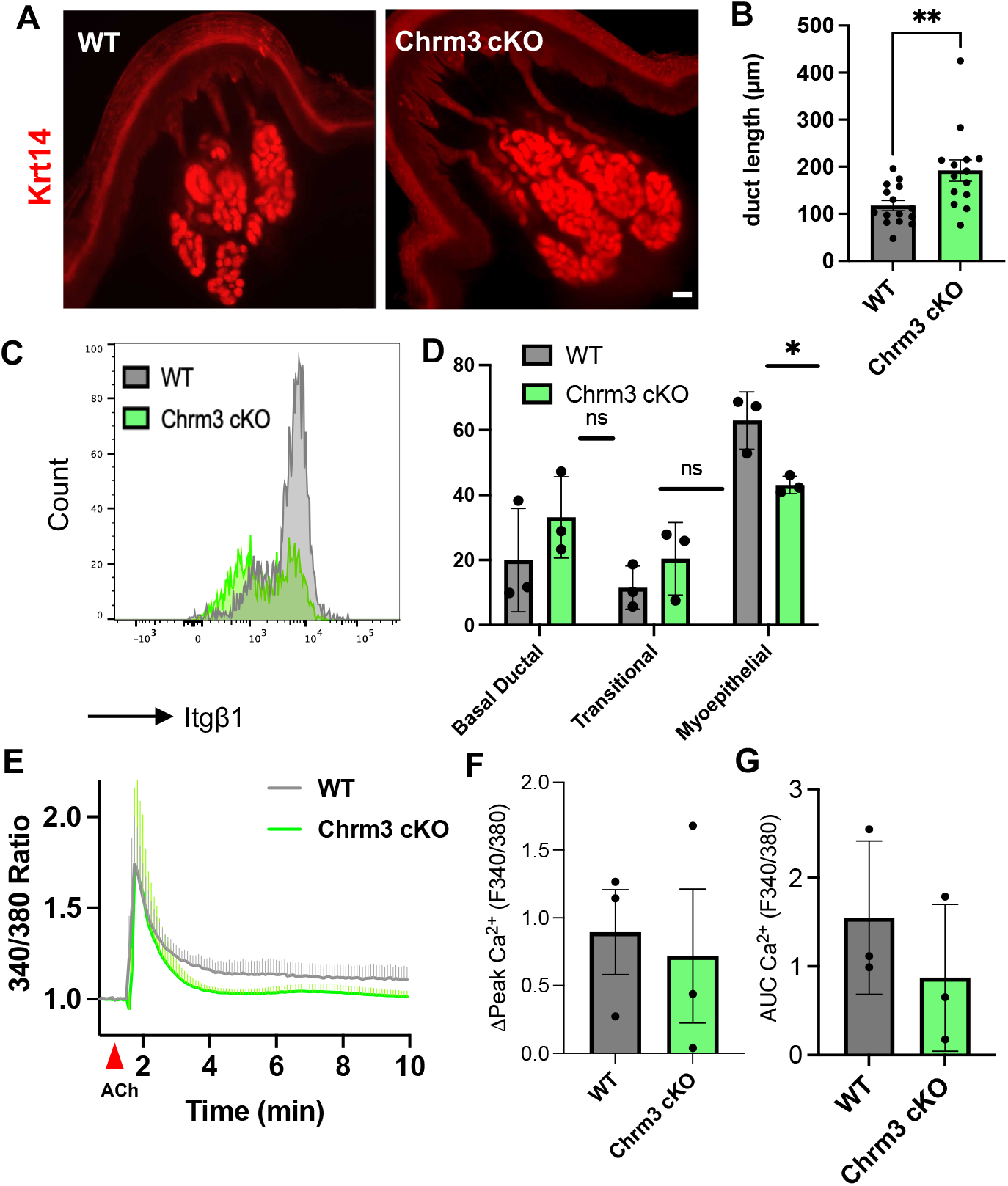
Cholinergic reception is important for glandular fate. **(A)** Representative slice from immunofluorescent image from whole-mount light-sheet imaging of WT and Chrm3 cKO sweat glands; keratin 14 (Krt14) keratinocyte marker **(B)** Quantification of duct length, unpaired t-test, p=0.02, 3 mice used per condition, each dot represents a measured ductal length **(C)** Representative FACS histogram of Itgβ1 expression in basal sweat gland populations in WT and Chrm3 cKO mice **(D)** Quantification of FACS analysis of cell populations present n=3 with 3 mice used in each sort per condition, unpaired t-test, p=0.01. Basal ductal= Sca-1^hi^ Itgβ1^lo^, Transitional= Sca-1^lo^ Itgβ1^lo^, Myoepithelial= Sca-1^lo^ Itgβ1^hi^ **(E)** Calcium imaging using the Fura-2 indicator of isolated sweat glands from WT and Chrm3 cKO mice with stimulus 1*μ*M acetylcholine (ACh) added at 1.5 min, traces represented as a mean of glands isolated from 2 WT and Chrm3 cKO mice (WT= 3 glands, Chrm3 KO= 3 glands), normalized baseline tracing before addition of stimulus to 1 **(F)** Quantification of peak response to ACh (highest value after stimulus addition-baseline calcium before stimulus) and **(G)** area under the curve (AUC) each dot represents an individual gland from 2 mice used in each condition, ns between groups

These results altogether demonstrate that both types of neurotransmitters are involved but at different stages of the sweat gland morphogenesis: NE through Adrb2 is important for sweat duct specification and ACh through Chrm3 is important for sweat gland differentiation.

### Loss of neuronal signals during development leads to upregulation of luminal cell features in basal cells

To explore the molecular mechanisms underlying neuronal signaling in sweat gland development, we acquired transcriptomic data from FACS-purified sweat duct basal cells and sweat gland myoepithelial cells from control and denervated mouse foot skin. The differentially expressed genes from ductal basal cells and glandular myoepithelial cells are shown in **Figure 5A-B**. Amongst these upregulated genes in denervated foot skin, we found a significant portion of them were shared by both myoepithelial cells and basal ductal cells **(Figure 5C, and Sup Table 1)**. Interestingly, many of the shared genes that are upregulated in both denervated basal cells are significantly involved in ion transport and glycolysis, and are normally expressed at a high level in luminal cells **(Figure 5D-E, S5)**. We further confirmed this finding using immunofluorescent imaging to demonstrate the ectopic expression of the luminal cell marker (Aqp5) in the myoepithelial cells (**Figure 5F**). Our results support the notion that neuronal signaling during development is critical for specification of basal cell fate, and that without neuronal signaling, both the ductal and glandular basal cells upregulate genes that normally define luminal cell fate.

**Figure 5.**
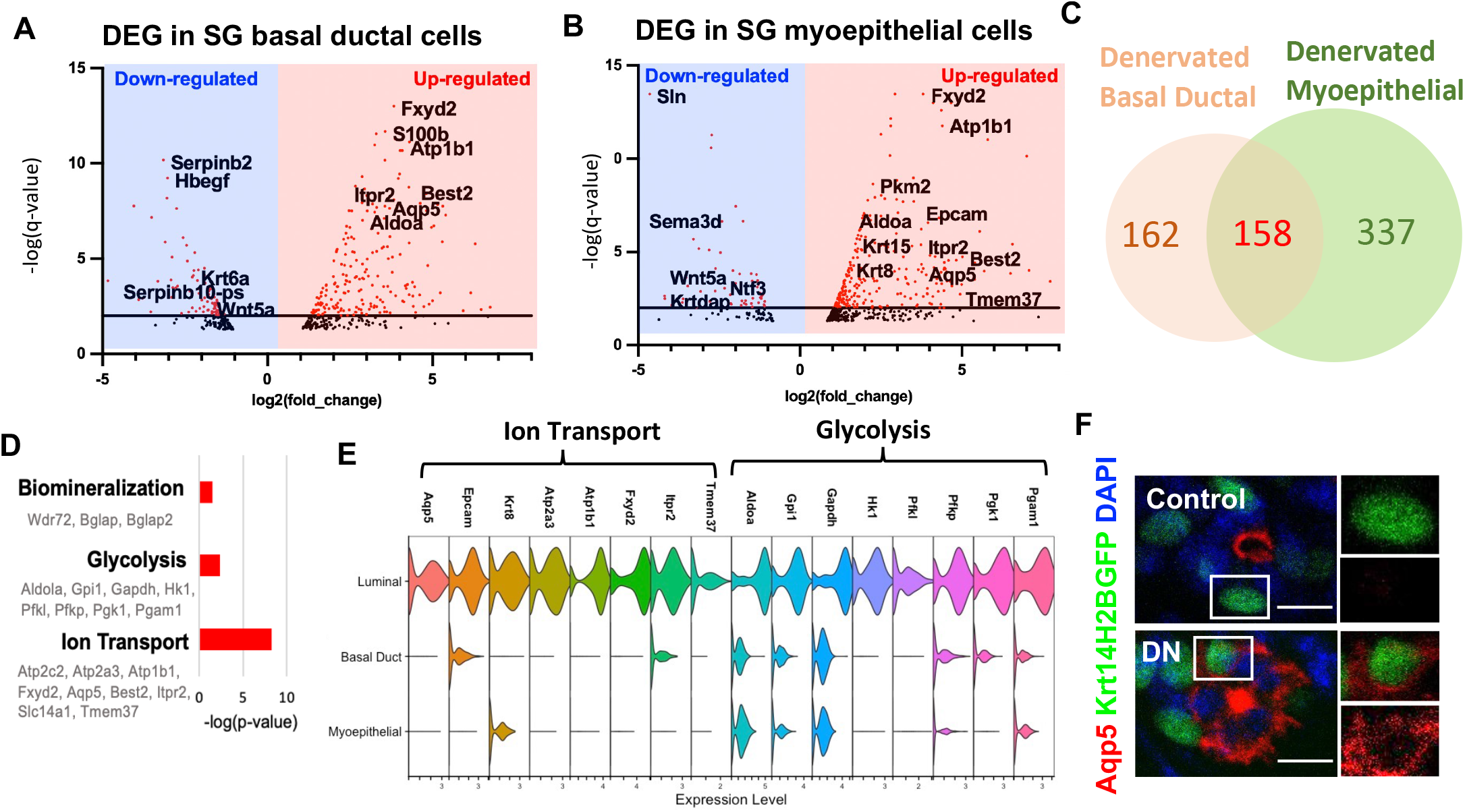
Loss of neuronal signals during development leads to upregulation of luminal cell features in basal cells. **(A)** Volcano plot of differentially expressed genes from bulk RNA-sequencing of FACS purified myoepithelial cells from adult mice that were denervated during development and **(B)** basal ductal cells **(C)** Venn diagram showing shared up-regulated genes in denervated myoepithelial and basal ductal cells **(D)** Pathway analysis of shared 158 up-regulated genes between myoepithelial and basal ductal cells **(E)** Violin plot of gene expression from WT single-cell RNA **(F)** Immunofluorescent imaging of sweat glands from control and denervated (DN) adult mice with various luminal cell markers with spilt channels for boxed are to right, Scale bars 10uM.

### Intracellular Ca^2+^ regulation by SLN is critical to establish sweat gland myoepithelial cell fate

Amongst the downregulated genes in myoepithelial cells upon denervation, sarcolipin (SLN) was the most significant candidate that was affected by loss of neuronal signaling (**Figure 5B**). Since SLN has been shown to be a critical Ca^2+^ regulator in cardiomyocytes and muscle cells (17-18), we sought to understand whether SLN is involved in sweat gland development through regulating intracellular Ca^2+^ levels. We analyzed the SLN knockout mice and first found that they exhibit an attenuated sweating response (**Figure S6A**); however, similar to denervated sweat glands, myoepithelial cells from SLN KO also showed ectopic expression of Aqp5 (**Figure 6A**). Further, we demonstrated that sweat glands from SLN KO mice showed decreased cytosolic Ca^2+^ upon ACh stimulation, similar to that in denervated and Chrm3 cKO sweat glands (**Figure 6B-D**).

**Figure 6.**
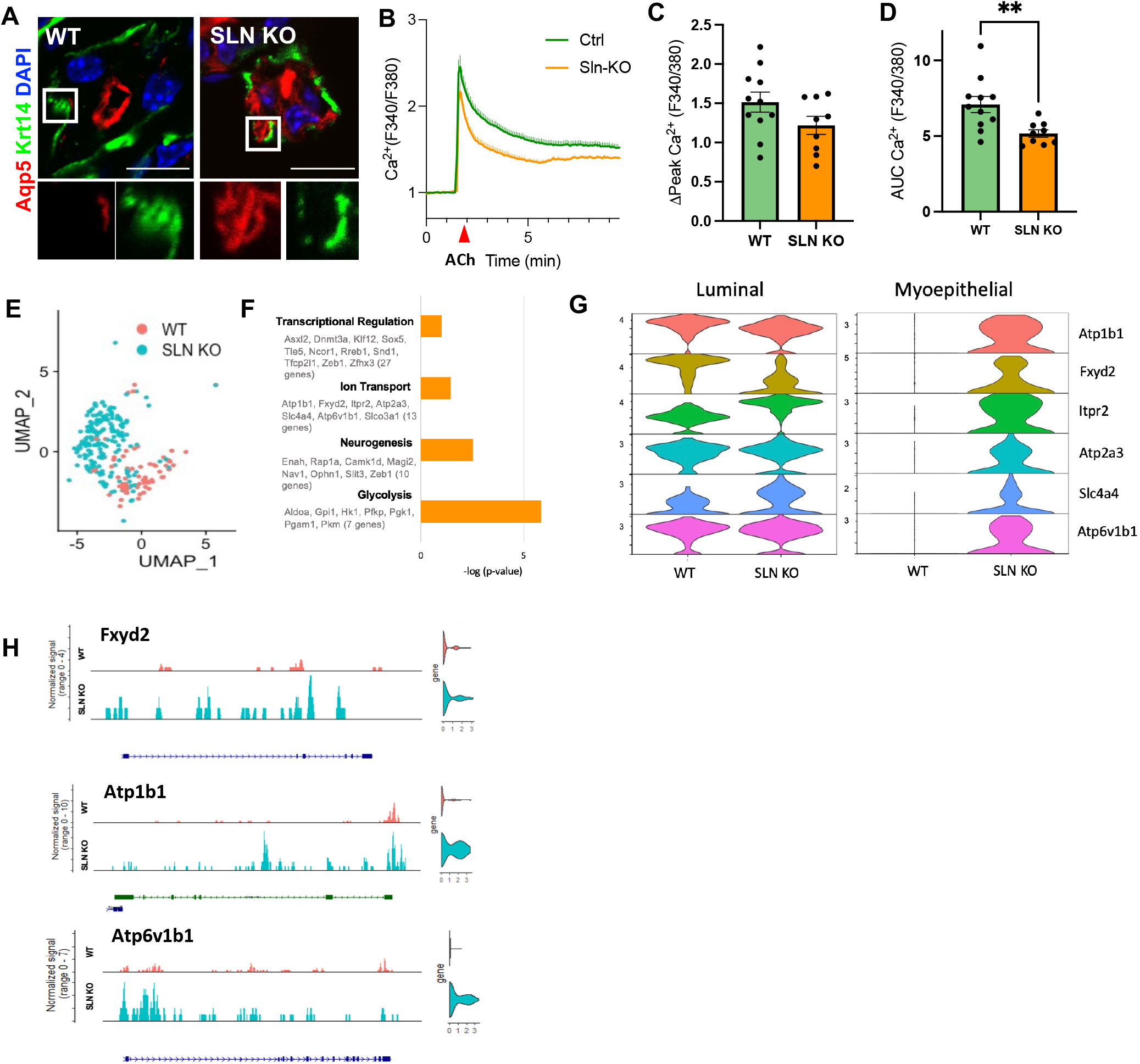
Intracellular Ca2+ regulation by SLN is critical to establish sweat gland myoepithelial cell fate. **(A)** Immunofluorescent imaging of sweat glands from WT and SLN KO adult mice with the luminal marker Aqp5 (Aquaporin 5). **(B)** Calcium imaging using the Fura-2 indicator of isolated sweat glands from WT and SLN KO mice with stimulus 1*μ*M acetylcholine (ACh) added at 1.5 min, traces represented as a mean of glands isolated from 4 WT and SLN KO mice (WT= 11 glands SLN KO= 9 glands), normalized baseline tracing before addition of stimulus to 1 **((C)** Quantification of peak calcium response and **(D)** area under curve (AUC) n=4 mice used in each condition, each dot represents and individual sweat gland’s response to ACh, unpaired t-test, p=0.0265 **(E)** UMAP projection of single-cell RNA-seq from WT and SLN KO myoepithelial cells (Acta2 >0.5) **(F)** Pathway analysis of gene changes in WT vs SLN KO sweat gland myoepithelial cells **(G)** Expression of luminal genes in luminal cells and myoepithelial cells **(H)** Single-cell ATAC-seq data of luminal cell genes in myoepithelial cells

To investigate the changes in the transcriptional landscape that may affect cell fate, we performed combined single cell RNA and ATAC sequencing for the foot skin samples from WT and SLN KO mice (**Figure 6E**). The myoepithelial cells from WT and SLN KO mice had different gene expression profiles (**Figure S6B**). Interestingly again, we found that many of the upregulated DEGs in SLN KO myoepithelial cells are involved in ion transport and glycolysis (**Figure 6F-G**). More specifically, genes such as Atp1b1, Fxyd2, Itpr2 and Atp6v1b1 that were exclusively expressed in the wildtype luminal cells are now expressed in the SLN myoepithelial cells (**Figure 6G**). We also found that there are a greater number of peaks in the chromatin regions of these genes in the SLN KO myoepithelial cells (**Figure 6H**), together suggesting that lower intracellular Ca^2+^ level upon the loss of SLN results in the upregulation of luminal cell genes in the myoepithelial cells.

### Sweat gland cell fate is specified by intracellular calcium level over time through reception of neuronal signals

As SLN KO sweat glands showed decreased cytosolic Ca^2+^ response upon ACh stimulation, accompanied with the ectopic expression of luminal cell genes, we hypothesized that high cytosolic Ca^2+^ is critical in the maintenance of glandular basal cell fate and that lowering cytosolic Ca^2+^ may lead to upregulation of the luminal cell features. To test this hypothesis, we isolated sweat duct progenitor cells from P3 mouse foot skin, and cultured in media with high or low Ca^2+^ to examine whether intracellular Ca^2+^ level affect the gene expression that specify glandular cell fate by immunofluorescent staining of Krt8 (luminal cell marker) and SMA (myoepithelial cell marker). We showed that sweat duct progenitor cells express higher level of Krt8 when cultured in low Ca^2+^, and higher level of SMA when cultured in high Ca^2+^, supporting the notion that high Ca^2+^ is critical in maintaining basal cell features while low Ca^2+^ triggers the expression of luminal cell features (**Figure 7A-B**).

**Figure 7.**
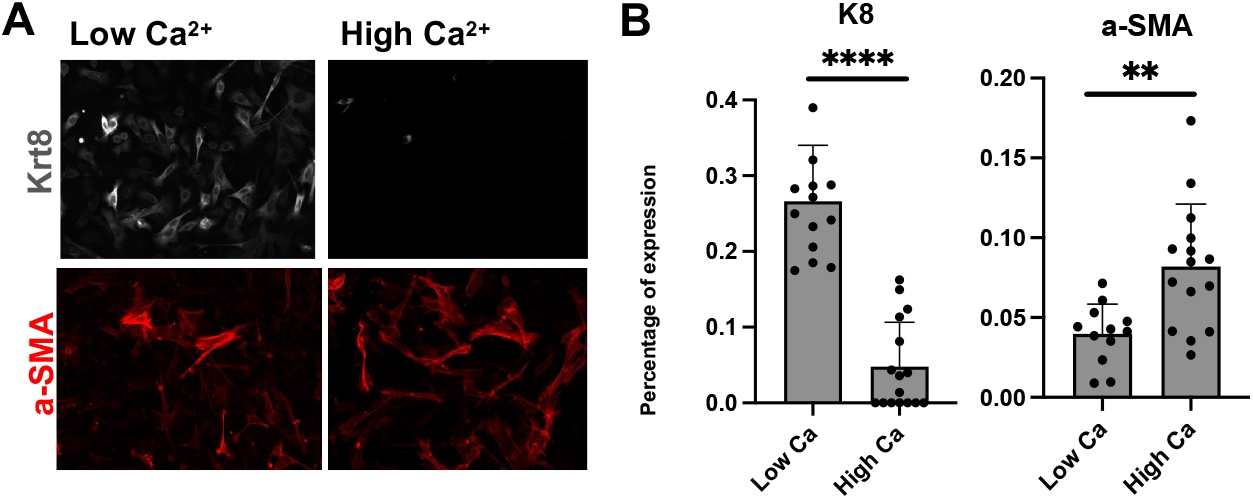
Sweat gland cell fate is specified by intracellular calcium level over time through reception of neuronal signals. **(A)** Imaging of culture with keratin 8 (Krt8; luminal cell marker) and smooth muscle actin (a-SMA; myoepithelial cell marker) in low calcium (50uM) and high calcium (3.6mM) **(B)** Quantification of percentage of cells meeting a defined cut-off value of expression, p=<0.0001, p=0.0022

## DISCUSSION

Our work provides a novel mechanism by which autonomic nerves impact tissue development, in addition to controlling the body’s physiologic responses. The sweat gland innervation is an intriguing system to study the role of autonomic nerves as they had been classically described undergoing a neurotransmitter switch from NE to ACh during nerve-gland co-development. This switch has remained puzzling as it was unclear if this switch had any functional role. In this work, we demonstrated that ductal cells express the Adrb2 receptor and respond to NE, while the glandular cells express the Chrm3 receptor and respond to ACh. The sequence of the neurotransmitter switch (NE then ACh) correlates with the appearance of ductal cells (Adrb2+) and later myoepithelial cells (Chrm3+) during morphogenesis, and we showed that ductal and glandular development is defective in the absence of the respective receptors. Therefore, our work provides a possible functional explanation for the switch as this switch is important for guiding the development of different parts of the sweat gland apparatus.

It is known that calcium is important in differentiation of cells and changes in calcium levels can impact gene expression via Ca^2+^-sensitive transcription factors, but it is not clear how the intracellular Ca^2+^ dynamics can exert long-term effects on cell fate specification or if intracellular Ca^2+^ levels fluctuate due to reception of neurotransmitters impacts the differentiation status of glandular keratinocytes.

We found that expression of Sarcolipin (SLN), a key regulator of sarcoendoplasmic reticulum (SR) Ca^2+^-ATPase (SERCA), is significantly down-regulated in the sweat gland myoepithelial cells upon denervation. The role of SLN has not been studied in tissues other than heart and muscle and it has not been shown that SLN expression can be controlled by neuronal signaling. Thus our work provides a novel role for a calcium regulator within the cell, SLN, that is altered with neuronal signaling and important in glandular development. We showed that intracellular Ca^2+^ level is important to specify the glandular cell fate and can be regulated beyond transient response upon the reception of neurotransmitters. We identified that Sarcolipin (SLN) expression in sweat gland myoepithelial cells is critical to regulate and maintain the high Ca^2+^ level as well as the basal cell fate, and its expression is lost in the absence of neuronal input during glandular development which leads to lower Ca^2+^ level and upregulation of the luminal cell fate. We also show a novel paradigm shift in the sweat gland. In culture, we showed that contrary to epidermal keratinocyte differentiation in which elevation of Ca^2+^ promotes epidermal differentiation (19), specification of the glandular myoepithelial (basal) cells requires high Ca^2+^ while lowering Ca^2+^ level promotes luminal (suprabasal) cell fate.

Previous work has shown how autonomic nerves can impact morphogenesis/differentiation in various target organs; however, these studies did not look at how mechanistically autonomic nerves are able to impact cell fate. In this work, we show an exciting novel link between neuronal signaling and calcium signaling and that calcium is a critical factor downstream of neuronal signals that regulates cell fate.

## Materials and Methods

### Mouse lines

C57BL/6 mice (Jax no. 000664), CD1 mice, K14H2BGFP and K14Cre provided by E. Fuchs (Rockefeller University) (20)(21), Adrb2^flox^ provided by G. Karsenty (Columbia University) (22), Chrm3^flox kindly^ provided by J. Wess (NIH) (23), SLN KO provided by G. Babu (Rutgers University) (17), GCAMP6f (Jax no. 028865)

### 6-OHDA Injection

Pups were injected once daily intraperitoneally with 6-hydroxydopamine hydrochloride (6-OHDA) 50ug/g of body weight (Sigma H4381) from post-natal day 0 to 5. 6-OHDA was made freshly and dissolved in 0.2% ascorbic acid PBS solution. Method adapted from Angeletti P. et al. 1971 (24).

### Sweat Test

Iodine-starch method- a 2% iodine/alcohol solution was applied to the paws of a restrained mouse and once the alcohol evaporated a mix of 1 g starch/1 ml castor oil was applied. Black dots appeared within 2 min on the footpad and finger-tips indicating active sweat pores (25).

### Immunohistochemistry

Mouse skin samples were fixed with 4% paraformaldehyde (PFA) for 15 min at room temperature and washed with PBS and left overnight in 30% sucrose at 4C (7). Samples were embedded in OCT compound. Sections of ∼10um were fixed in PFA for 5 min and washed with PBS x3, washed in 0.1% Triton X-100 in PBS for 15 min, blocked in blocking buffer (1% BSA, 1% Gelatin, 0.25% Normal Goat Serum, 0.25% Normal Donkey Serum, 0.30% TritonX-100, 1X PBS) for 1 hour at RT then stained with primary antibodies over night at 4C. Slides were then washed with PBS x3 and stained with secondary antibody for 2 hours at RT. Antibodies and dilutions are as follows: Smooth muscle actin (rabbit, 1:200, Abcam), Keratin 14 (chicken, 1:800, BioLegend), Integrin-Beta 1 (rat, 1:100, Alexa647-conjugated, BioLegend), Aquaporin 5 (rabbit, 1:1500, Abcam), Keratin 8 (rabbit, 1:800, Abcam), EpCAM (rat, 1:100, Invitrogen), Sca-1 (rat, 1:100 Alexa488-conjugated, BioLegend),Tyrosine Hydroxylase (sheep, 1:200, Millipore). Secondaries and dilutions are as follows: Donkey anti-rabbit Cy3 (1:400, Jackson ImmunoResearch), Donkey anti-rabbit A488 (1:800, Invitrogen), Donkey anti-sheep 488 (1:400, Abcam), donkey anti-chicken Alexa Fluor 488 (1:800, Jackson ImmunoResearch), donkey anti-chicken Alexa Fluor 647 (1:400, Jackson ImmunoResearch), Rhodamine Red Donkey anti-sheep (1:200 Jackson ImmunoResearch). The slides were then washed with 1X PBS x3. Coverslips were then applied with Prolong Gold antifade reagent (Invitrogen).

### Image acquisition and processing

Images were acquired using a Zeiss 880 confocal and Zeiss AxioObserver through a 20x air and 63x oil objective. RGB images were assembled using Fiji software.

### FACS

Mouse paw pads were collected as biopsies and incubated in 50mM EDTA, 37°C, for 30 min After manual dissection of epidermis, the dermal fraction containing sweat glands was placed in Hanks Balanced Salt Solution (Lonza) with 1000U/ml collagenase, 300U/ml hyaluronidase (Sigma), 1g/1uL Liberase TL (Roche), and digested for 1.5hr at 37°C. 0.25% Trypsin-EDTA (Gibco) was added for additional digestion for 10min. Digested tissues were suspended and washed with PBS containing 4% of fetal bovine serum (FBS), then filtered through 40μm cell strainers to make single cell suspensions. Samples were labeled with fluorescently-conjugated antibodies for 30 min at 4C-Sca1-APCCy7 (1:1500, eBioscience), α6-PE (1:1000, eBioscience) and α6-FITC (1:200, Invitrogen), β1-Alexa647 (1:500, eBioscience) at 4°C for 30min. Then washed with PBS/FBS. DAPI was added for dead cell exclusion. Cell sorting was performed on SY300 cell sorter and FACSAria IIu sorter and analysis was performed using FlowJo software.

### Sweat gland isolation

Footpads and tips of mouse digits were dissected under a stereoscope microscope and digested for 45 min in 1g liberase TL (Roche)/1mL HBSS with calcium at 37C. A wide-orifice 1000 uL pipette was used to release the sweat glands. The solution with sweat glands was transferred to an Eppendorf tube and spun down at 300xG for 3 min. The gland tissue was then used for calcium measurements.

### Calcium Imaging

Measurement of intracellular calcium in isolated sweat glands was done by a technique developed by Concepcion et al. 2016 (26). Sweat glands were seeded onto Cell-Tak (Corning) coated coverslips and loaded with 1uM Fura-2-AM (Invitrogen) and 0.5 uM Cell Tracker Orange (Invitrogen) for 30 min. Calcium was measured using a time-lapse imaging on a IX81 epifluorescence microscope (Olympus). Sweat glands were perfused with 2mM Ca^2+^ ringer solution (5M NaCl, 1M KCl, 1M CaCl_2_, 1M MgCl_2_, 1M Hepes, D-Glucose) and stimulated with 1uM acetylcholine and 1uM norepinephrine. A ROI was draw on each sweat gland and the ratio of Fura-2 emission following excitation at 340 nm (F340) and 380 nm (F380) were calculated for each time point and analyzed using SlideBook imaging software v4.2 (Olympus).

### Bulk RNA sequencing of myoepithelial cells and basal ductal cells

FACS purified samples were submitted to the Genomics Core Laboratory of Weill-Cornell Medical College for RNA extraction, labeling, and hybridization to Affymetrix Mouse 430A_2 arrays. Statistical analyses were performed using R version 2.13.0, Bioconductor version 2.6 and Spotfire Decision Site 9.1.1 (Tibco, Somerville, MA). Gene lists were uploaded to the “Database for Annotation, Visualization and Integrated Discovery” (DAVID) v2022q4web tool to assess pathway enrichment.

### Single-cell RNA sequencing

P0, P3, P6, and adult mouse foot skin was digested using the digestion protocol for FACS. Cell hashing was used to label samples from distinct samples (27). Single cell suspensions were labeled with the TotalSeq Hashtag Antibodies (BioLegend) for 30 min on ice. Single cell suspensions were submitted to the Genome Technology Core facility at New York University Langone Medical Center and subjected to 10X genomics sequencing. Analysis of the data was performed using the Seurat V3 package (Stuart and Butler et al. Comprehensive Integration of Single-Cell Data. Cell (2019) [Seurat V3]) (28). Quality control was performed to calculate the numbers of genes and UMIs for each cell. Cells with a low number of covered genes (gene-count <200) were filtered out. Cells that were negative for the hashtag labeling or a doublet were filtered out so the data only included singlets. The data was normalized, scaled, and variable features were used to perform PCA. The FindNeighbros and FindClustes functions were then used and we then performed Uniform Manifold Approximation and Projection (UMAP) to visualize the data set. The developmental time-points (P0, P3, P6) were integrated with the adult keratinocytes by first selecting all Krt14 positive cells and then merging the datasets, splitting the dataset in two Seurat objects, normalizing and identifying the variable features for each dataset independently. The features that are repeatable variable across datasets were selected for integration. The command IntegrateData was then used to create an integrated data assay. The data was scaled, PCA was performed, the FindNeighbros and FindClustes functions were used and we then performed UMAP to visualize the data set.

### Single-cell RNA/ATAC Sequencing

WT, 6-OHDA treated, SLN KO adult mouse foot skin was digested using the digestion protocol for FACS. Cells were FACS sorted to enrich for keratinocytes. Nuclei isolation was performed using the 10X Genomic protocol for low cell input nuclei isolation for single cell multiome ATAC + gene expression sequencing. Single cell suspensions were submitted to the Genome Technology Core facility at New York University Langone Medical Center and subjected to multiome RNA-ATAC sequencing. Analysis of the data was performed using the Seurat V3 package (Stuart and Butler et al. Comprehensive Integration of Single-Cell Data. Cell (2019) [Seurat V3]) (28) and Signac 1.9.0 package (Stuart et al. Single-cell chromatin state analysis with Signac. Nature Methods (2021) (29). RNA and ATAC data was loaded and an object containing the RNA data was created. An ATAC assay was created and added to the object. Quality control was performed by removing cells with low and unusually high counts for both the RNA and ATAC assay. Gene expression was normalized, and dimensionality was reduced using PCA. Latent semantic indexing was performed (LSI) for the scATAC data. Weighted nearest neighbor method was used for joint UMAP visualization. Data was integrated as in the above single-cell sequencing method. Gene lists were uploaded to the “Database for Annotation, Visualization and Integrated Discovery” (DAVID) v2022q4web tool to assess pathway enrichment.

### Whole-Mount imaging

A Zeiss Z.1 light sheet microscope was used for whole-mount imaging at 2X. Tissue was cleared using the CUBIC method (Susaki et al. 2015) (30). Tissue was stained with primary antibody for three days and secondary antibody for three days in blocking buffer solution after CUBIC-1 step. Images were assembled in Imaris 9.8.0. Duct length quantification was performed by measuring the straight ductal region with measure tool.

### Calcium culture

Progenitor cells were isolated from post-natal day 3 pups and cultured. E-media (with low Ca^2+^ (50 uM) and high Ca^2+^ (3.6 mM) and 15% serum conditions was added for one day. After one day, the cells were fixed with 4%PFA/PBS for 15 min and stained according to staining protocol in immunohistochemistry section. Images were quantified by selection of each cell and creating a threshold intensity and quantifying the number of cells with signal above the established threshold.

### Antibodies

Sca1-APCCy7 (1:1500, BD, 560654)

α6-PE (1:1000, eBioscience, 555734)

α6-FITC (1:200, Invitrogen, 11-0495-82),

β1-Alexa647 (1:500, BioLegend, 102214)

Smooth muscle actin (rabbit, 1:200, Abcam)

Keratin 14 (chicken, 1:800, BioLegend, 906001)

Aquaporin 5 (rabbit, 1:1500, Abcam, ab78486)

Keratin 8 (rabbit, 1:800, Abcam, ab59400)

EpCAM (rat, 1:100, eBioscience, 11-5791-82)

Sca-1 (rat, 1:100 Alexa488-conjugated, BioLegend, 122516)

Tyrosine Hydroxylase (sheep, 1:200, Millipore, AB1542)

Donkey anti-rabbit Cy3 (1:400, Jackson ImmunoResearch, 711-165-152)

Donkey anti-rabbit A488 (1:800, Jackson ImmunoResearch, 711-545-152)

Donkey anti-sheep 488 (1:400, Abcam, ab150177)

Donkey anti-chicken Alexa Fluor 488 (1:800, Jackson ImmunoResearch, 703-545-155)

Donkey anti-chicken Alexa Fluor 647 (1:400, Jackson ImmunoResearch, 703-605-155)

Rhodamine Red Donkey anti-sheep (1:200 Jackson ImmunoResearch),

DAPI Solution (Fisher Scientific, 1:3000, D1306)

### Chemicals and Other Reagents

HBSS (ThermoFisher, 2402117)

Collagenase (Sigma, C2674)

Hyaluronidase (Sigma, H3506-500MG)

Trypsin/EDTA (Corning, MT25053CI)

Liberase TL (Sigma, 5401020001)

Cell-Tak (Corning, 354240)

6-Hydroxydopamine hydrochloride (6-OHDA) (Sigma, H4381)

Acetylcholine chloride (Sigma, A6625)

DL-Norepinephrine hydrochloride (Sigma, A7256)

Fetal Bovine Serum

Fura-2 (ThermoFisher, F1221)

Cell-tracker (ThermoFisher C34551)

### Software and Algorithms

R R Development Core Team 2022 http://www.R-project.org

Seurat CRAN https://satijalab.org/seurat/

Signac https://stuartlab.org/signac/index.html

DAVID Knowledgebase v2022q4 https://david.ncifcrf.gov/

## Acknowledgements

We thank facilities at NYU Langone: Cytometry and Cell Sorting Laboratory (P30CA016087), Experimental Pathology Research Laboratory (S10 OD021747), the Applied Bioinformatics Laboratories, the Genome Technology Center (P30CA016087), and NYU Langone’s Microscopy Laboratory (P30CA016087, S10RR023704). We would also like to thank Edrogmus S., for assistance with calcium imaging experiments, Freidman C., Manahil N., and Omene B. for preliminary data processing. C.L. and lab members are supported by start-up fund from Hansjörg Wyss Department of Plastic Surgery. Research reported in this publication was supported by *the Eunice Kennedy Shriver National Institute of Child Health and Human Development* of the National Institutes of Health under award number 1F30HD105455-01. The content is solely the responsibility of the authors and does not necessarily represent the official views of the National Institutes of Health.

## Author contributions

J.R. and C.P.L. conceived the project and designed experiments. J.R. and J.T. performed experiments and analyses. M.L. provided advanced computational analyses and assisted in image whole-mount image acquisition. A.C. and S.F. assisted in calcium imaging experiments and shared their protocol for imaging and analysis and provided feedback/expertise on calcium signaling and manuscript preparation. S.M. and G.B. provided guidance in experiments relating to SLN. J.R. and C.L. wrote the manuscript.

## References

1. LeBouef T, Yaker Z, Whited L. Physiology, Autonomic Nervous System. 2022 May 8. In: StatPearls [Internet]. Treasure Island (FL): StatPearls Publishing; 2022 Jan–. PMID: 30860751.

2. Ceasrine AM, Lin EE, Lumelsky DN, Iyer R, Kuruvilla R. Adrb2 controls glucose homeostasis by developmental regulation of pancreatic islet vasculature. Elife. 2018 Oct 10;7:e39689. doi: 10.7554/eLife.39689. PMID: 30303066; PMCID: PMC6200393

3. Kamiya A, Hayama Y, Kato S, Shimomura A, Shimomura T, Irie K, Kaneko R, Yanagawa Y, Kobayashi K, Ochiya T. Genetic manipulation of autonomic nerve fiber innervation and activity and its effect on breast cancer progression. Nat Neurosci. 2019 Aug;22(8):1289–1305. doi: 10.1038/s41593-019-0430-3. Epub 2019 Jul 8. PMID: 31285612.

4. Knox SM, Lombaert IM, Haddox CL, Abrams SR, Cotrim A, Wilson AJ, Hoffman MP. Parasympathetic stimulation improves epithelial organ regeneration. Nat Commun. 2013;4:1494. doi: 10.1038/ncomms2493. PMID: 23422662; PMCID: PMC3582394.

5. Knox SM, Lombaert IM, Reed X, Vitale-Cross L, Gutkind JS, Hoffman MP. Parasympathetic innervation maintains epithelial progenitor cells during salivary organogenesis. Science. 2010 Sep 24;329(5999):1645–7. doi: 10.1126/science.1192046. PMID: 20929848; PMCID: PMC3376907.

6. Magnon C, Hall SJ, Lin J, Xue X, Gerber L, Freedland SJ, Frenette PS. Autonomic nerve development contributes to prostate cancer progression. Science. 2013 Jul 12;341(6142):1236361. doi: 10.1126/science.1236361. PMID: 23846904.

7. Zhang B, Ma S, Rachmin I, He M, Baral P, Choi S, Gonçalves WA, Shwartz Y, Fast EM, Su Y, Zon LI, Regev A, Buenrostro JD, Cunha TM, Chiu IM, Fisher DE, Hsu YC. Hyperactivation of sympathetic nerves drives depletion of melanocyte stem cells. Nature. 2020 Jan;577(7792):676–681. doi: 10.1038/s41586-020-1935-3. Epub 2020 Jan 22. PMID: 31969699; PMCID: PMC7184936.

8. Shwartz Y, Gonzalez-Celeiro M, Chen CL, Pasolli HA, Sheu SH, Fan SM, Shamsi F, Assaad S, Lin ET, Zhang B, Tsai PC, He M, Tseng YH, Lin SJ, Hsu YC. Cell Types Promoting Goosebumps Form a Niche to Regulate Hair Follicle Stem Cells. Cell. 2020 Aug 6;182(3):578-593.e19. doi: 10.1016/j.cell.2020.06.031. Epub 2020 Jul 16. PMID: 32679029; PMCID: PMC7540726.

9. Landis SC, Keefe D. Evidence for neurotransmitter plasticity in vivo: developmental changes in properties of cholinergic sympathetic neurons. Dev Biol. 1983;98(2):349–372.

10. Yodlowski ML, Fredieu JR, Landis SC. Neonatal 6-hydroxydopamine treatment eliminates cholinergic sympathetic innervation and induces sensory sprouting in rat sweat glands. J Neurosci. 1984;4(6):1535-1548.1984

11. Guidry G, Landis SC. Sympathetic axons pathfind successfully in the absence of target. J Neurosci. 1995 Nov;15(11):7565–74. doi: 10.1523/JNEUROSCI.15-11-07565.1995. PMID: 7472507; PMCID: PMC6578085.

12. Schotzinger RJ, Landis SC. Cholinergic phenotype developed by noradrenergic sympathetic neurons after innervation of a novel cholinergic target in vivo. Nature. 1988 Oct 13;335(6191):637–9.

13. Schotzinger RJ, Landis SC. Acquisition of cholinergic and peptidergic properties by sympathetic innervation of rat sweat glands requires interaction with normal target. Neuron. 1990 Jul;5(1):91

14. Lin MJ, Lu CP. Glandular stem cells in the skin during development, homeostasis, wound repair and regeneration. Exp Dermatol. 2021 Apr;30(4):598–604. doi: 10.1111/exd.14319. Epub 2021 Mar 22.

15. Lu CP, et al. Identification of stem cell populations in sweat glands and ducts reveals roles in homeostasis and wound repair. Cell. 2012;150:136–150.

16. Cui CY, Schlessinger D. Eccrine sweat gland development and sweat secretion. Exp Dermatol. 2015;24:644–650.

17. Babu GJ, Bhupathy P, Timofeyev V, Petrashevskaya NN, Reiser PJ, Chiamvimonvat N, Periasamy M. Ablation of sarcolipin enhances sarcoplasmic reticulum calcium transport and atrial contractility. Proc Natl Acad Sci U S A. 2007 Nov 6;104(45):17867–72. doi: 10.1073/pnas.0707722104. Epub 2007 Oct 30. PMID: 17971438; PMCID: PMC2077025.

18. Pant M, Bal NC, Periasamy M. Sarcolipin: A Key Thermogenic and Metabolic Regulator in Skeletal Muscle. Trends Endocrinol Metab. 2016 Dec;27(12):881–892. doi: 10.1016/j.tem.2016.08.006. Epub 2016 Sep 13.

19. Bikle DD, Xie Z, Tu CL. Calcium regulation of keratinocyte differentiation. Expert Rev Endocrinol Metab. 2012 Jul;7(4):461–472

20. Tumbar T, Guasch G, Greco V, Blanpain C, Lowry WE, Rendl M, Fuchs E. Defining the epithelial stem cell niche in skin. Science. 2004 Jan 16;303(5656):359–63. doi: 10.1126/science.1092436. Epub 2003 Dec 11. PMID: 14671312; PMCID: PMC2405920.

21. Vasioukhin V, Degenstein L, Wise B, Fuchs E. The magical touch: genome targeting in epidermal stem cells induced by tamoxifen application to mouse skin. Proc Natl Acad Sci U S A. 1999 Jul 20;96(15):8551–6. doi: 10.1073/pnas.96.15.8551. PMID: 10411913; PMCID: PMC17554.

22. Hinoi E, Gao N, Jung DY, Yadav V, Yoshizawa T, Myers MG Jr, Chua SC Jr, Kim JK, Kaestner KH, Karsenty G. The sympathetic tone mediates leptin’s inhibition of insulin secretion by modulating osteocalcin bioactivity. J Cell Biol. 2008 Dec 29;183(7):1235–42. doi: 10.1083/jcb.200809113. Epub 2008 Dec 22. PMID: 19103808; PMCID: PMC2606962.

23. Gautam D, Han SJ, Hamdan FF, Jeon J, Li B, Li JH, Cui Y, Mears D, Lu H, Deng C, Heard T, Wess J. A critical role for beta cell M3 muscarinic acetylcholine receptors in regulating insulin release and blood glucose homeostasis in vivo. Cell Metab. 2006 Jun;3(6):449–61. doi: 10.1016/j.cmet.2006.04.009. PMID: 16753580.

24. Angeletti, PU, Lev-Montalcini R. Chemical sympathectomy in newborn animals. 1971. Neuropharmacology 10:55–59

25. Sato K, Sato F. Methods for studying eccrine sweat gland function in vivo and in vitro. Methods Enzymol. 1990;192:583–99.

26. Concepcion AR, Vaeth M, Wagner LE 2nd, Eckstein M, Hecht L, Yang J, Crottes D, Seidl M, Shin HP, Weidinger C, Cameron S, Turvey SE, Issekutz T, Meyts I, Lacruz RS, Cuk M, Yule DI, Feske S. Store-operated Ca2+ entry regulates Ca2+-activated chloride channels and eccrine sweat gland function. J Clin Invest. 2016 Nov 1;126(11):4303–4318.

27. Stoeckius M, Zheng S, Houck-Loomis B, Hao S, Yeung BZ, Mauck WM 3rd, Smibert P, Satija R. Cell Hashing with barcoded antibodies enables multiplexing and doublet detection for single cell genomics. Genome Biol. 2018 Dec 19;19(1):224.

28. Stuart T, Butler A, Hoffman P, Hafemeister C, Papalexi E, Mauck WM 3rd, Hao Y, Stoeckius M, Smibert P, Satija R. Comprehensive Integration of Single-Cell Data. Cell. 2019 Jun 13;177(7):1888-1902.e21.

29. Stuart T, Srivastava A, Madad S, Lareau CA, Satija R. Single-cell chromatin state analysis with Signac. Nat Methods. 2021 Nov;18(11):1333–1341.

30. Susaki EA, Tainaka K, Perrin D, Yukinaga H, Kuno A, Ueda HR. Advanced CUBIC protocols for whole-brain and whole-body clearing and imaging. Nat Protoc. 2015 Nov;10(11):1709–27.

