## Supplemental Figures for "Neurotransmitter signaling specifies sweat gland stem cell fate through SLN-mediated intracellular calcium regulation"

### Supplementary Figure 1

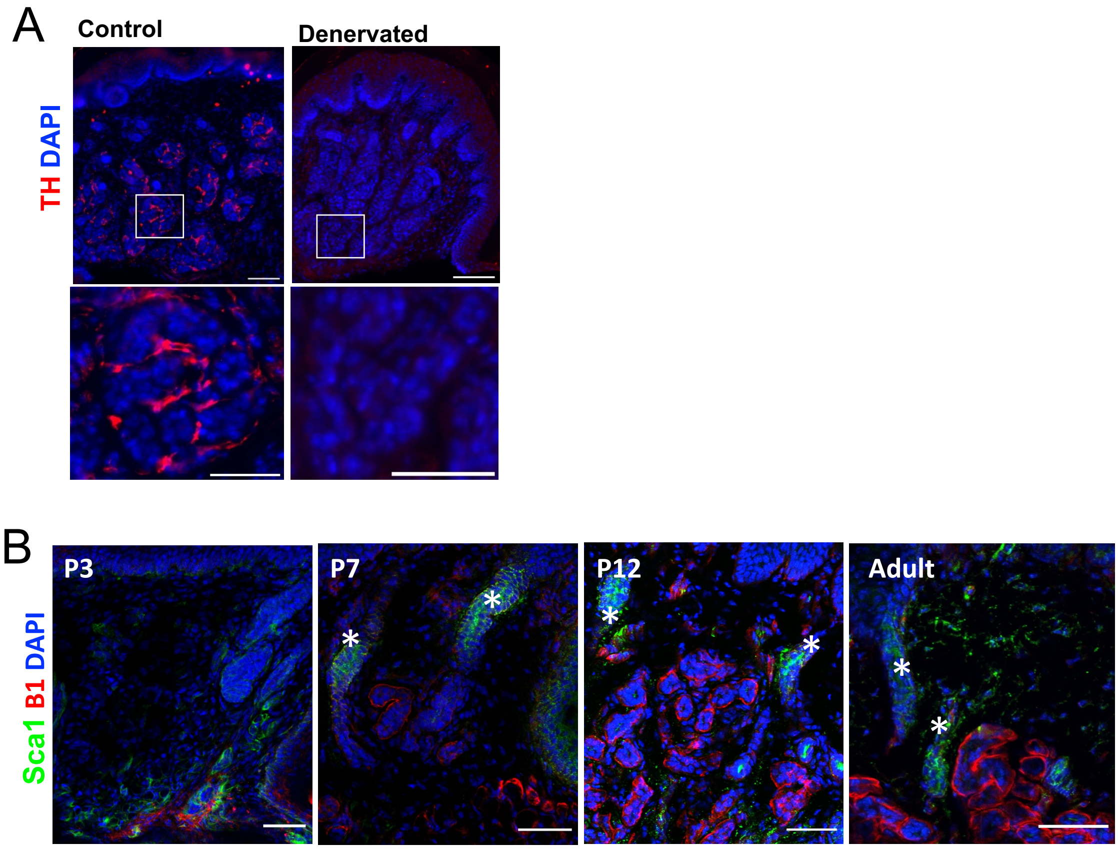


**(A)** Immunofluorescent imaging of sweat glands from mice injected with PBS (control) or 6-OHDA (denervated) from P0-P5 at 6 months of age. Scale bars 100uM large image, 50uM mag image (**B)** Immunofluorescent imaging of sweat glands from post-natal day 3 (P3) 7, 12, and adult mice. * indicates duct. Scale bars 50uM.

### Supplementary Figure 2

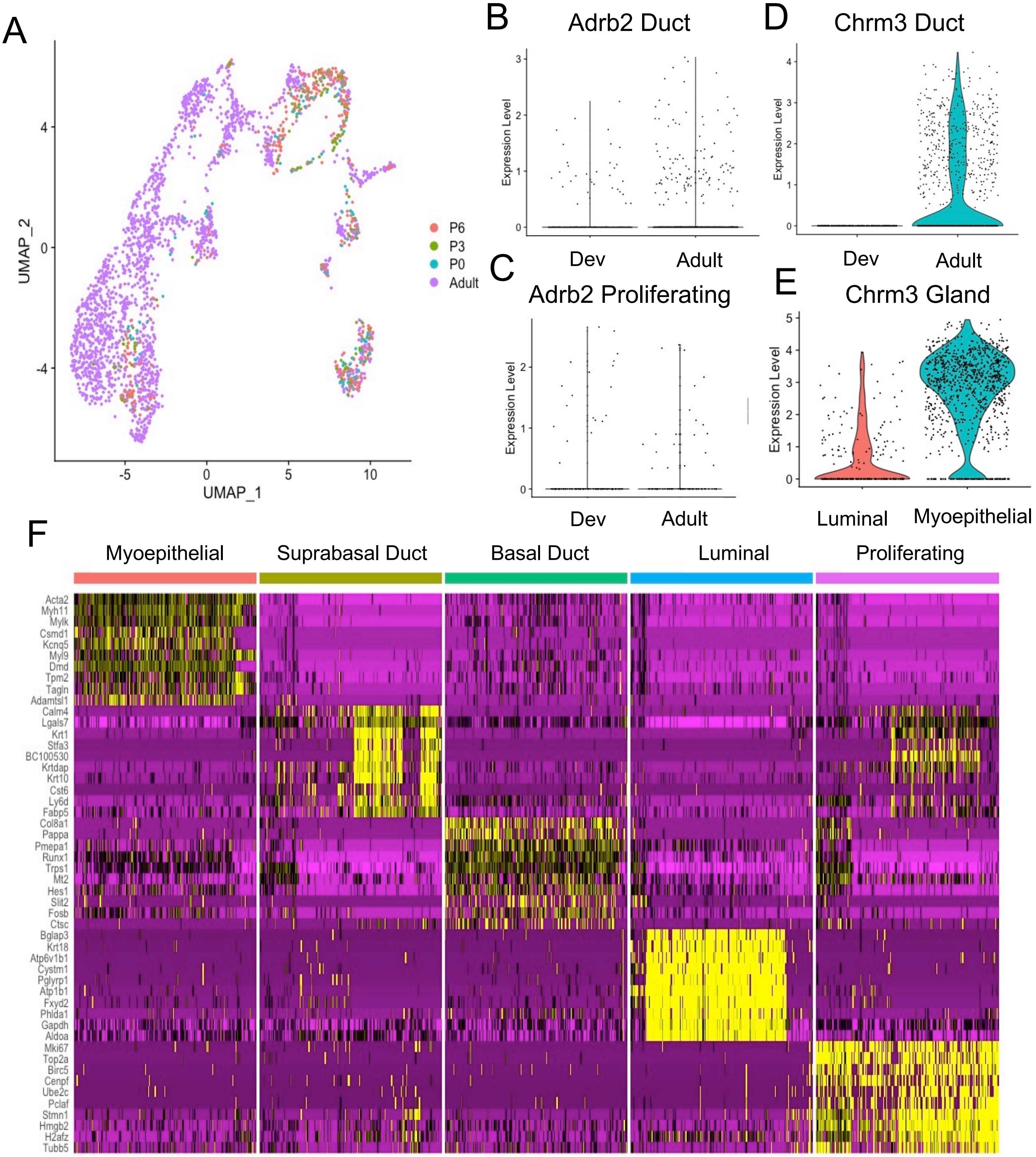


**(A)** UMAP projection of single-cell RNA-seq (scRNA-seq) data from sweat glands of P0, P3, P6, and adult mice spilt by developmental stage **(B)** Violin plot of Adrb2 receptor expression in ductal clusters (suprabasal ductal and basal ductal) in developmental (dev) and adult **(C)** Violin plot of Adrb2 receptor expression in proliferating cluster (Ki67+) in developmental (dev) and adult **(D)** Violin plot of Chrm3 receptor expression in ductal clusters (suprabasal ductal and basal ductal) in developmental (dev) and adult **(E)** Violin plot of Chrm3 expression in glandular luminal and myoepithelial cells **(F)** Heat map showing top genes in each cluster of single-cell RNA sequencing data

### Supplementary Figure 5

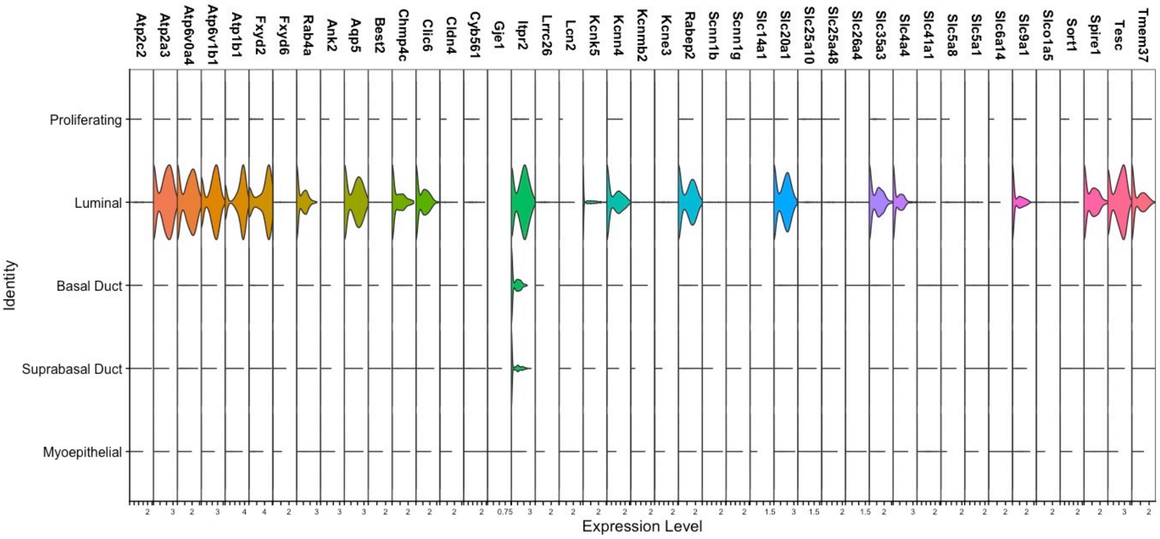


Violin plot using single-cell RNA-sequencing from control sweat gland keratinocytes of ion transport genes

### Supplementary Figure 6

A

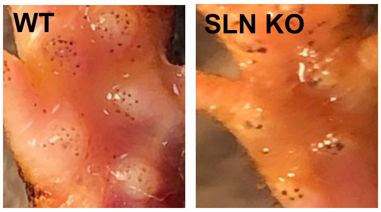


B

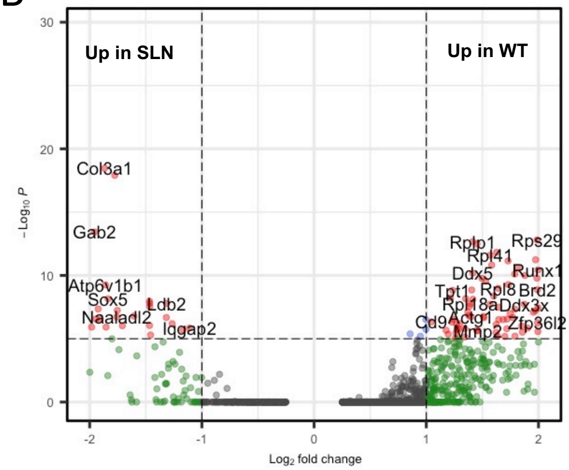


**(A)** Iodine sweat test of natural sweating response from WT and SLN KO adult mice **(B)** Volcano plot from single-cell RNA sequencing of WT and SLN KO myoepithelial cells

### Supplementary Table 1

| Gene | Denervated Duct log2(fold_change) | Denervated Myoepithelial log2(fold_change) |
|---|---|---|
| Atp2c2 | 3.64422 | 3.644 |
| Atp2a3 | 4.36054 | 4.36054 |
| Atp6v0a4 | 3.03146 | 2.91776 |
| Atp6v1b1 | 5.58258 | 5.58258 |
| Fxyd2 | 3.82697 | 3.827 |
| Fxyd6 | 1.34361 | 2.63317 |
| Rab4a | 2.34282 | 1.23331 |
| Ank2 | 3.64763 | 3.25845 |
| Aqp5 | 3.56208 | 3.76642 |
| Best2 | 4.28428 | 4.28428 |
| Chmp4c | 2.22569 | 3.1829 |
| Clic6 | 5.82822 | 4.52192 |
| Cldn4 | 2.88099 | 4.99232 |
| Cyb561 | 2.23798 | 3.0658 |
| Gje1 | 10.8084 | 6.56809 |
| Hbb-b2 | 2.27319 | 1.63926 |
| Itpr2 | 2.946 | 3.73435 |
| Lrrc26 | 3.47091 | 4.44083 |
| Lcn2 | 2.21979 | 2.80013 |
| Kcnk5 | 3.18527 | 4.46497 |
| Kcnn4 | 4.19358 | 4.94419 |
| Kcnmb2 | 6.48344 | 9.88838 |
| Kcne3 | 3.78126 | 4.44113 |
| Rabep2 | 2.44409 | 2.70898 |
| Scnn1b | 3.02656 | 4.17395 |
| Scnn1g | 2.29406 | 1.99107 |
| Slc14a1 | 4.00216 | 3.98743 |
| Slc20a1 | 2.86801 | 2.31524 |
| Slc25a10 | 1.42847 | 1.73713 |
| Slc25a48 | 3.54918 | 4.03213 |
| Slc26a2 | 1.16058 | 1.58948 |
| Slc26a4 | 6.16082 | 8.88027 |
| Slc35a3 | 1.33244 | 1.77648 |
| Slc4a4 | 2.96393 | 3.6058 |
| Slc41a1 | 1.27169 | 1.07777 |
| Slc5a1 | 1.48944 | 4.35994 |
| Slc6a14 | 4.00216 | 1.07777 |
| Slc9a1 | 1.79241 | 1.42767 |
| Slco1a5 | 2.3563 | 4.14819 |
| Sort1 | 1.71851 | 2.91684 |
| Spire1 | 2.6952 | 2.26555 |
| Tesc | 4.62918 | 5.40468 |
| Tmem37 | 4.29104 | 4.69794 |

Table of the fold change from control in denervated basal ductal and myoepithelial cells of the shared up-regulated genes from bulk sequencing of FACS isolated cells
